# VirtualIce: Half-synthetic CryoEM Micrograph Generator

**DOI:** 10.1101/2024.09.28.615520

**Authors:** Alex J. Noble

## Abstract

Single particle cryo-electron microscopy (cryoEM) is going through a phase of rapid optimization focused on increasing the efficiency, accuracy, and automation of every step in the data pipeline. Machine learning models in particular have been making substantial advances in cryoEM, however their impact has been limited. This limitation is due in part to the lack of availability of realistic ground-truth datasets for training and evaluation of cryoEM machine learning models. To address this limitation and accelerate this phase, we introduce VirtualIce which generates half-synthetic micrographs by projecting proteins onto real, curated micrographs of vitrified buffer. VirtualIce provides configurable features including noise simulation, realistic particle distributions, particle overlapping, particle aggregation, filtering obscured regions, and multiple structures per micrograph. VirtualIce may be a valuable resource to help visualize unknown proteins, accelerate the development of automated data collection and processing pipelines, and develop cryoEM algorithms.

## Introduction

Cryo-electron microscopy (cryoEM) allows for high-resolution studies of structures of biological molecules^1,2^. Over the past decade, cryoEM has emerged from a niche technique to a critical component in structural biology research, comparable to x-ray crystallography in the number of structure releases^3,4^. CryoEM algorithm and pipeline development for data collection and processing has accelerated in the direction of automation^5–9^ and intelligent data selection and filtering^9–15^. However, several hurdles remain in the development of highly optimized, automated workflows, including optimal targeting during collection, micrograph selection based on particle distribution, particle picking, particle angular assignment, particle classification, heterogeneity analysis, and resolution estimation and prediction. The more recent emergence of effective machine learning methods has amplified opportunities to overcome these hurdles, provided that enough sufficiently realistic ground-truth data exists for training and evaluation.

Here we present VirtualIce, a customizable open-source Python tool that generates half-synthetic cryoEM micrographs, addressing the need for ground-truths and enabling researchers to visualize how their proteins might appear in cryoEM. While existing software and methods focus on the forward model for generating micrographs^16–21^ or only simulate particle projections^8,22–25^, VirtualIce focuses on creating realistic micrographs using real vitrified buffer micrographs and simulating realistic particle behaviors in the ice (note: cryoET simulators exist as well^26–34^). We are additionally releasing the vitrified buffer dataset with manually curated masked contamination and grid substrate areas for use as realistic backgrounds in VirtualIce-generated micrographs. VirtualIce uses the masks of these obscured regions to avoid saving particle coordinates in these areas, adding real context to each particle’s environment and enabling the creation of realistic ground-truth datasets.

VirtualIce use-cases that will be discussed include generating ground-truth data for developing improved methods for particle picking, heterogeneity analysis, classification, preferred orientation assessment, and resolution prediction, along with teaching cryoEM and simulating live pipelines for collection and processing. VirtualIce provides extensive customization to the user regarding particle behavior including particle distribution, aggregation, overlap, removal of obscured particle coordinates, and multiple structures per micrograph. VirtualIce can be called and wrapped by other software. VirtualIce source code is freely available under the MIT license (https://github.com/alexjnoble/VirtualIce) and the labeled vitrified buffer dataset is available in the public domain (EMPIAR-12287). Refer to the Ethical Use Agreement for proper use. VirtualIce requires EMAN2^25^ for some operations and benefits from an IMOD^35^ installation for visualization. VirtualIce runs on both multi-CPU and multi-GPU systems.

## Results

VirtualIce offers a simple command-line user interface for generating half-synthetic micrographs. Once the user has installed VirtualIce and downloaded the provided vitrified buffer micrographs and associated labels, the user may start generating micrographs by specifying structures (PDB IDs, EMDB IDs, and local .pdb/.mrc/.map files) as in the following example command:

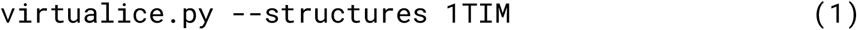

This command will generate micrographs based on many default settings by projecting a random number of randomly-oriented EM maps of PDB 1TIM (triose phosphate isomerase)^36^ onto randomly chosen vitrified buffer micrographs, distributed according to the relative ice thickness of the micrograph (proxied by low-resolution feature variations), potentially with overlapping particles and some particle aggregation, then save the micrographs as .mrc files and save the particles not overlapping with each other or with obscured regions to a Relion-compatible .star file. The projection process includes pre-orienting the mrc created from the pdb so that it’s major axis is along the y-axis of the micrograph before any protein orientation has been applied (useful for algorithm development), generating a projection from a Poisson distribution to simulate electron counts, convolving the projection with CTF based on the buffer micrograph using EMAN2, and adding Gaussian white noise based on the micrograph’s estimated noise parameters to a collage of particles before adding the collage to the micrograph. A micrograph generated from this command is shown in Figure 1a. Figure 1b-d show additional micrographs generated from PDBs [1DAT (apoferritin)^37^, 7ZP8 (ribosome)^38^, and 2HCO (hemoglobin)^39^], 1RUZ (hemagglutinin)^40^, and 1PMA (proteasome)^41^ with the following commands, respectively:

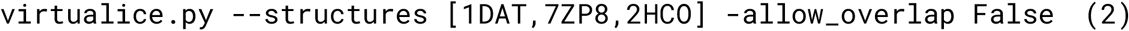

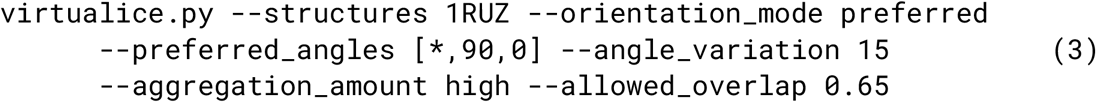

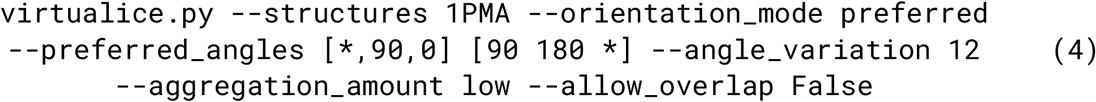

**Figure 1.**
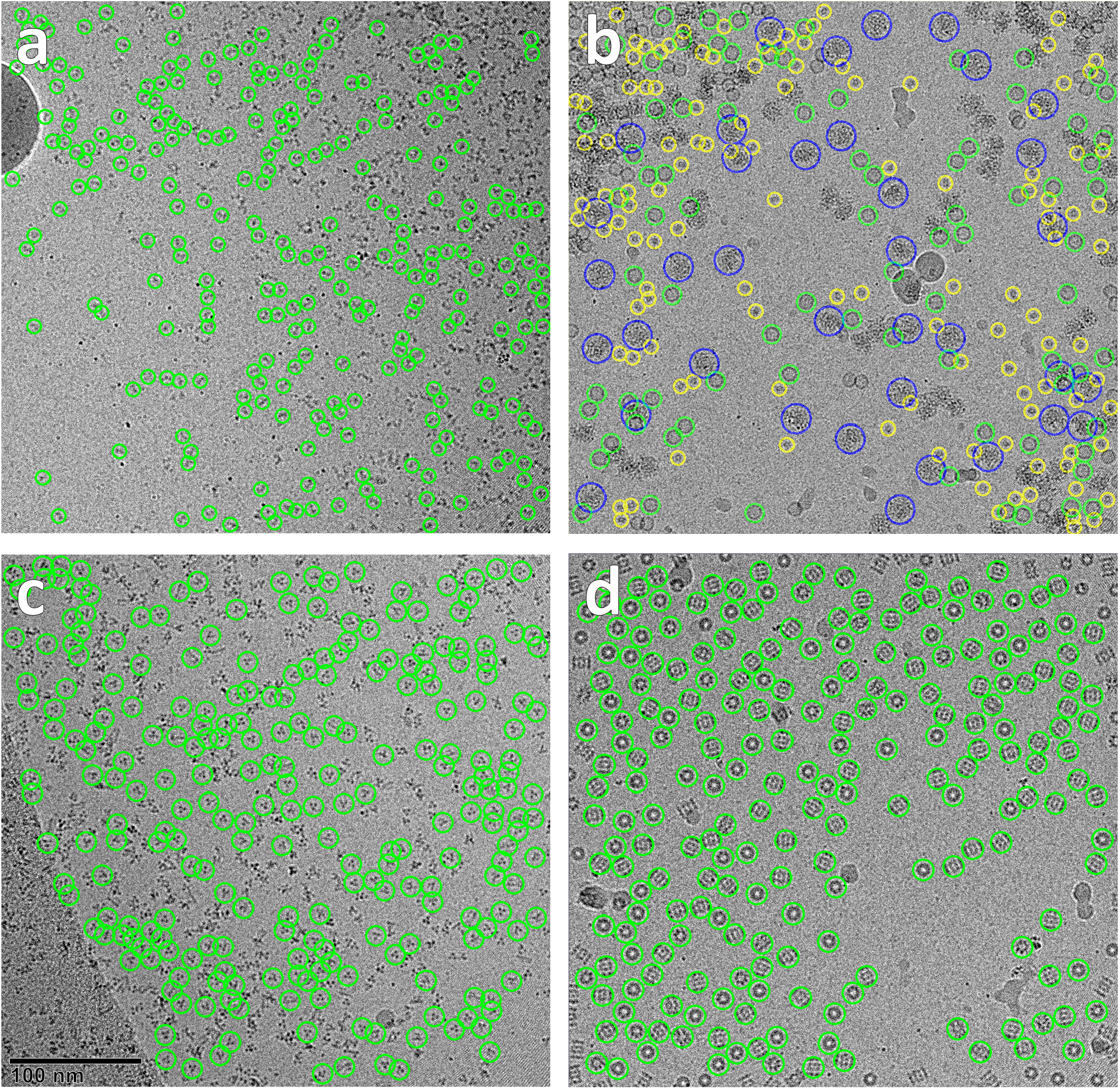
Assortment of VirtualIce-generated micrographs. A selection of micrographs generated with VirtualIce illustrating several features. (a) PDB 1TIM projected onto a vitrified buffer micrograph with non-overlapping particles saved to coordinate files (green circles). Ice contamination and particles too close to the edge are also avoided when saving coordinates. Particle distribution matches the apparent ice thickness. (b) PDBs 1DAT (green), 7ZP8 (blue), and 2HCO (yellow) projected together onto a buffer micrograph, exemplifying the multi-structure ability of VirtualIce. (c) PDB 1RUZ showing one preferred orientation and avoiding the carbon substrate. (d) PDB 1PMA showing two preferred orientations and not overlapping.

The micrographs in Figure 1 include labels encircling the centers of each particle, excluding particles overlapping or nearly overlapping with other particles, overlapping with obscured regions, and too close to the micrograph edge. Supplementary Figure 1 shows the micrographs in Figure 1 without labels. Saving coordinates that are not obscured, overlapping, or too close to the edge of the micrograph ensures that the particle picks are clean, providing ground-truth data for many machine learning applications. The user may also extract the particles at their full resolution with the `--crop_particles` flag, save coordinates as additional IMOD^35^ .mod or (x,y,z) .coord files, Fourier downsample the micrographs, and save micrographs additionally in PNG and/or JPEG formats, amongst other output options.

Figure 1a (and Fig S1a) shows a 53 kDa protein projected on top of a vitrified buffer micrograph where overlap is allowed. The particle distribution correlates with the apparent ice thickness of the vitrified buffer micrograph where there is a gradient of thicker to thinner ice from the top-right to the bottom-left; the particle density is higher in thicker ice. Ice contamination (including on the top-left and near the top-right) and beam fringes (top-right and bottom-right) present in the vitrified buffer micrograph contribute to the realistic context of a typical cryoEM micrograph. By default, particles saved to coordinate files (green circles) include only particles that are not overlapping with other particles, with obscuring objects in the vitrified buffer micrograph, and with particles too close to the edge of the micrograph. Figure 1b (and FigS1b) shows a multi-structure micrograph with apoferritin (442 kDa, green circles), ribosome (1385 kDa, blue), and hemoglobin (65 kDa, yellow) proteins where overlap is allowed. The same observations with respect to particle distribution, contamination, fringes, and saved particles as Figure 1 apply. Figure 1c (and FigS1c) shows hemagglutinin (165 kDa) where overlap is allowed. Here, hemagglutinin is restricted to orientations that closely match what is seen in plunge-frozen grids: most hemagglutinin proteins have strong preferred orientation where primarily their top-views are present and are allowed to tilt slightly relative to this preferred orientation. In addition to the observations made in Figures 1a,b, this vitrified micrograph includes carbon grid substrate, which VirtualIce avoids when saving coordinates. Figure 1d (and FigS1d) shows proteasome (686 kDa) where overlap is not allowed. Proteasome is almost entirely restricted to top and side views, mimicking its real behavior. The same observations as in Figure 1a,b are also present here.

VirtualIce gives users control over several particle and micrograph behavior features, including the amount of particle aggregation, ice thickness, and particle distribution, in addition to preferred orientation and particle overlapping shown in Figure 1. Figure 2 shows micrographs of PDB 8EIQ (CFTR; 173 kDa)^42^ with varying degrees of aggregation, ranging from 0 (none) to 10 (high), with the figure showing aggregation levels of (a) 0, (b) 5, and (c) 10. Figure 3 shows synthetic micrographs of PDB 8G02 (bacteriophage complex; 136 kDa)^43^ with increasingly thick simulated ice thickness, represented by protein contrast, of (a) 30, (b) 100, and (c) 200 nm. Figure 4 shows synthetic micrographs of PDB 6AHU (ribonuclease P; 493 kDa)^44^ with each particle distribution option available in VirtualIce. Figures 2-4 were created with the following command templates, respectively, where * signifies the parameter that was varied.

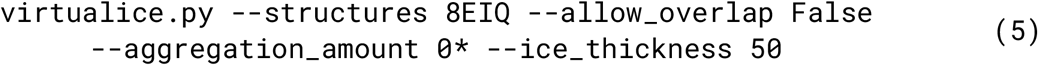

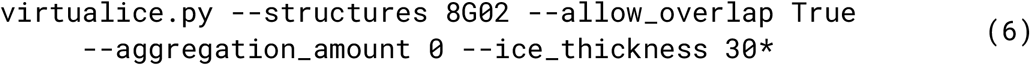

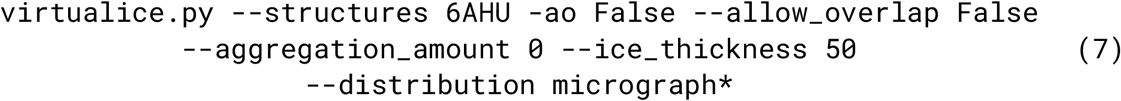

**Figure 2.**
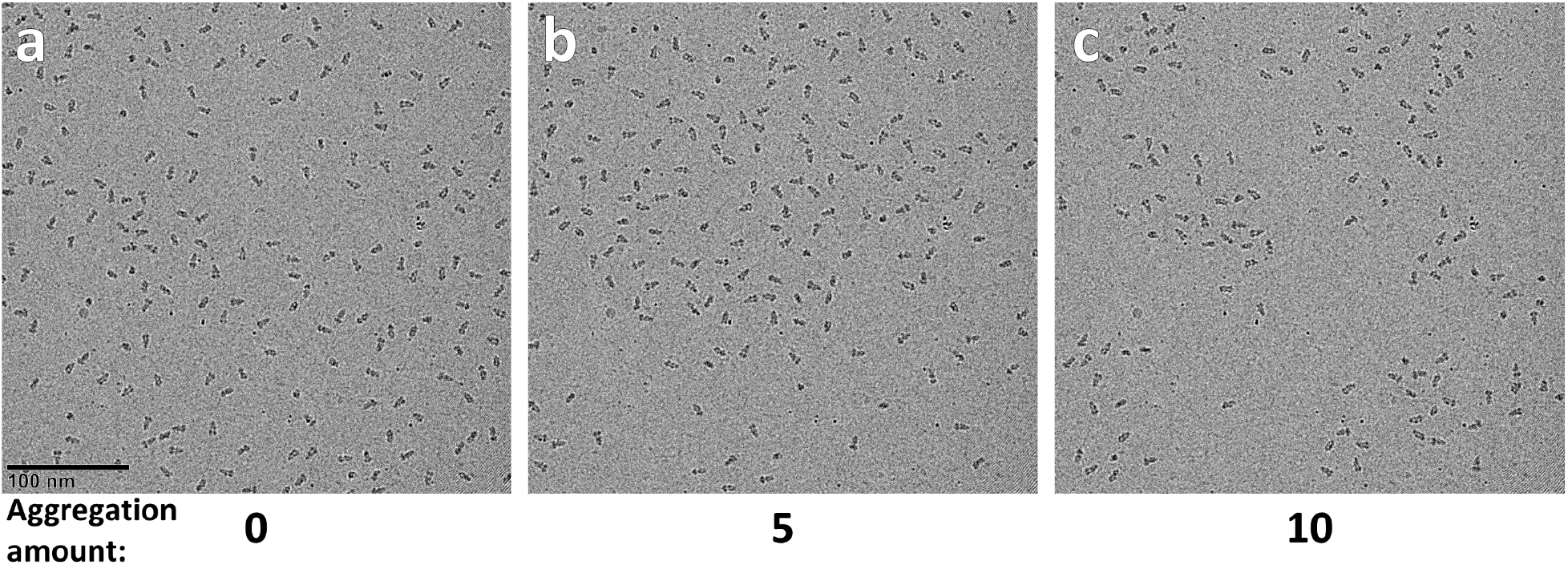
VirtualIce-generated micrographs exhibiting varying protein aggregation. (a) PDB 8EIQ projected onto a vitrified buffer micrograph with no overlapping particles and no aggregation. (b) Same as (a) but with 5/10 aggregation. (c) Same as (a) but with 10/10 aggregation.

**Figure 3.**
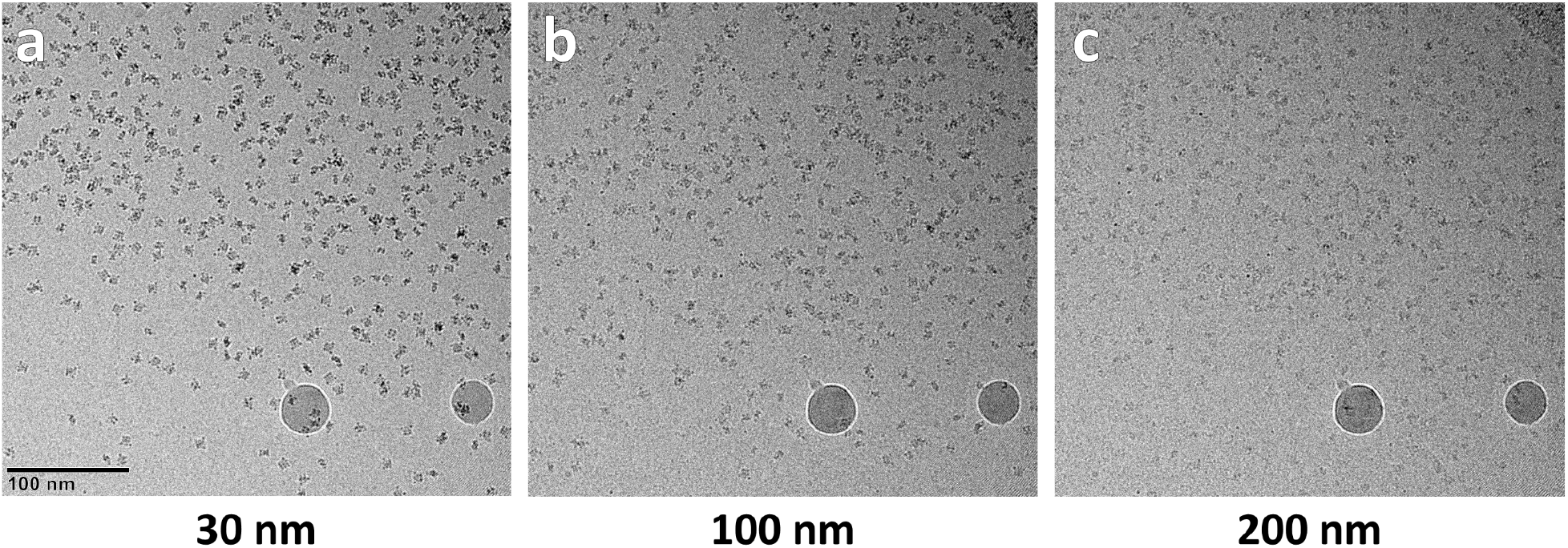
VirtualIce-generated micrographs displaying simulated ice thickness. (a) PDB 8G02 projected onto a vitrified buffer micrograph with a simulated ice thickness of 30 nm, reflected in the amount of particle contrast. (b) Same as (a) but with 100 nm simulated ice thickness. (c) Same as (a) but with 200 nm simulated ice thickness.

**Figure 4.**
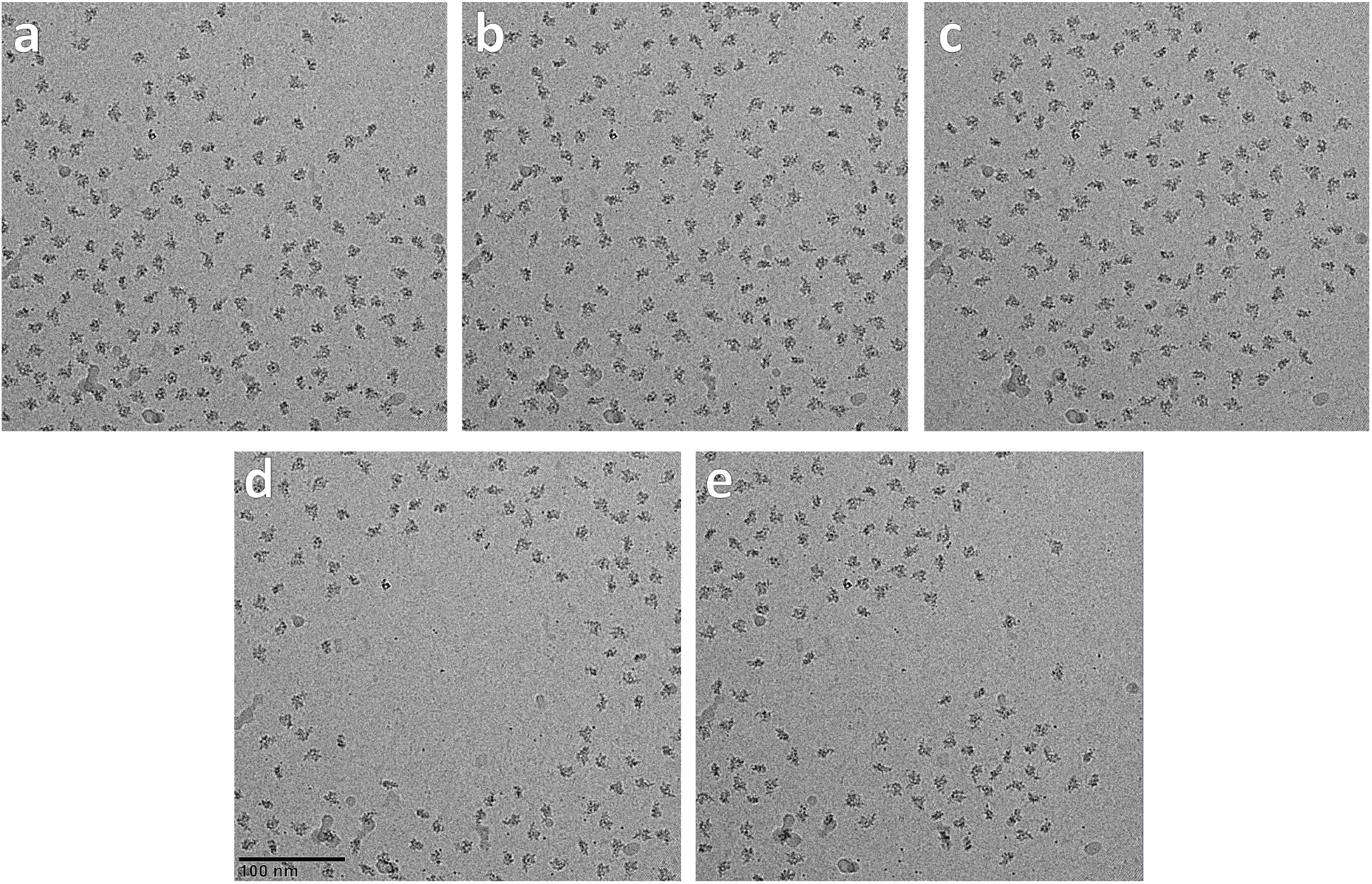
VirtualIce-generated micrographs showing particle distribution options. (a) PDB 8G02 projected onto a vitrified buffer micrograph using the ‘micrograph’ distribution option which places particles in darker areas of the micrograph (bottom-left) with higher probability than lighter areas (top-right), indirectly modeling the effect of ice thickness on particle distribution. (b-e) Same vitrified buffer micrograph as in (a) but with ‘random’, ‘circular’, ‘inverse circular’, and random ‘gaussian’ blob distributions, respectively.

## Methods

### Vitrified Buffer Micrographs

To maximize the realism of the generated micrographs, a dataset of vitrified buffer was collected across a range of clean ice and contaminated ice, both fully within cryoEM grid holes and with carbon substrate in some micrographs. The sample was prepared without protein using Spotiton^45^ and a carbon nanowire grid^46^. The grid had irregularly-dispersed gold flakes, visible at high magnification, due to contamination of preparation instruments from Quantifoil gold grids that used to have this problem. ∼900 micrographs were collected with Leginon^47^ on a Titan Krios 300 keV cryoTEM (Thermo Fisher Scientific) and Gatan K2 camera with a Gatan BioQuantum energy filter (15 eV slit width) (Gatan Inc. Pleasanton, CA, USA), a pixelsize of 1.096 Å, total dose of 50.81 e-/Å^2^, nominal defocus range randomly chosen between 1 and 3 µm, and spherical aberration near 0.0 due to a Cs corrector. 75 frames were collected for each micrograph, which were aligned with Motioncor2^48^ and the resulting micrograph’s CTF was estimated using CTFFind4^49^, both through Appion^50^.

### Manual Labeling of Vitrified Buffer Micrographs

AnyLabeling v0.3.3^51^ was used to manually label objects that can obscure particles projected with VirtualIce, including crystalline ice contamination, grid substrate, and contamination from gold flakes. Micrographs were first converted from grayscale to a green tint to reduce eye stress during labeling and to make objects such as grid substrate, ice contamination, and gold flakes stand out better (Fig. S2). The Mobile Segment Anything Model (MobileSAM)^52^ within AnyLabeling was used to assist with manual labeling, which was useful primarily for labeling ice contamination and substrate, but less useful for labeling gold flakes. 678 micrographs were manually labeled over a few months.

### VirtualIce Implementation

The VirtualIce workflow is versatile, allowing for many different types and forms of input and output. Here we describe every step in the VirtualIce workflow.

VirtualIce can generate single- and multi-structure micrographs using any protein in PDB or MRC/MAP formats. The `--structures` flag accepts as input any number of single or multiple structure sets, either as single entries (single-structure) or bracketed, comma/space-separated entries (multiple-structures), as exemplified in commands `(1)` - `(7)`. `--structures` also accepts `random` for a random PDB/EMDB download, `rp` for a random PDB download, `re` for a random EMDB download, and local .pdb/.mrc/.map files. The user specifies the number of micrographs to generate, `--num_images`, and optionally specifies the number of particles per micrograph, `--num_particles`, otherwise this will be random up to the maximum number of particles (< 100 particles is down-weighted) that can fit in the micrograph multiplied by the softly-conditioned number of particle layers, `--num_particle_layers`, if the `--allow_overlap` flag is set. VirtualIce downloads the symmetrized version of the requested PDB from RCSB^53^ if it exists, then converts it to an MRC using EMAN2^25^ given the pixelsize of the vitrified buffer micrographs (`--apix`) to scale properly and conversion resolution (`--pdb_to_mrc_resolution`). Imported or downloaded MRC/MAP files are normalized, scaled to the micrograph pixelsize using EMAN2, and thresholded to try to remove dust in the EM map (`--std_threshold`). PDB files that were converted to MRC and optionally imported or downloaded MRC/MAP files are re-oriented so that the longest axis points in the z-direction, which are projected onto micrograph so that the z-axis of the protein volume aligns with the y-axis of the micrograph before any rotation of the protein. The protein mass is estimated, using either EMAN2 for PDB files or an internal algorithm for imported/downloaded MRC/MAP files, which is recorded to an information file that VirtualIce produces. Using VirtualIce with PDB files is recommended because PDB conversions to MRC produce EM maps without noise or dust around the protein, which standardizes thresholding as opposed to imported/downloaded MRC/MAP files, thus more reliably producing realistic half-synthetic micrographs.

Each micrograph is generated by first randomly selecting a vitrified buffer micrograph (`--image_directory`) with its associated defocus value stored in a text file (`--image_list_file`). The protein aggregation amount is selected based on user input (`--aggregation_amount`), particle overlap is determined randomly if not specified by the user (`--allow_overlap`), and noise parameters of the selected vitrified buffer micrograph are estimated for use later. The minimum cube-size of the protein MRC is determined and the MRC is padded up to an FFT-friendly box size (`--scale_percent`, default 33.33% larger) to allow for more delocalized CTF information to be present from each particle. A user-specified or randomly-determined ice thickness value (`--min_ice_thickness` and `--min_ice_thickness` or `--ice_thickness`) is converted to an empirically-determined fudge-factor for adding particles to the micrograph at a later step. A user-specified 2D particle distribution is used to place particles in the micrograph (`--distribution`; micrograph: mimics vitrified buffer micrographs’ local ice thickness, probabilistically placing more particles in darker areas, gaussian: random number of gaussian particle clumps, circular: placed in a circular area, inverse_circular: placed outside a circular area, random: randomly placed based on a uniform distribution). The micrograph distribution option is recommended for producing the most realistic-looking micrographs. The user may decide whether particles can go to the edge of the micrograph (`--no_edge_particles`) and by how much (`--border`).

Protein orientation is determined before proteins are projected. By default, particles will be randomly oriented, but the user may instead specify that the particles be uniformly oriented or preferentially-oriented with user specified orientations (`--orientation_mode`). Preferred orientations are user-specified (`--preferred_angles`) by inputting any number of three Euler angles (ZYZ rotation convention), where wildcards mean that the protein may rotate freely along that axis (e.g. [*,90,0] means the protein is free to rotate along the first z-axis, then rotates 90 degrees around the y-axis, and is not rotated around the new z-axis). Equal sampling will be applied to each preferred orientation in the user-specified list. Preferentially-oriented particles are then perturbed by an additional random angle within a small range (`--angle_variation`) to mimic real preferred orientation variations due to proteins not being completely relaxed after being adsorbed to the air-water interface right before vitrification^54^. When preferred orientations are requested, only a percent (default 90%) of proteins will be preferentially oriented while the rest will be randomly oriented (`--preferred_weight`), mimicking the percentage of proteins in typical cryoEM samples that are adsorbed to the air-water interface and thus preferentially-oriented^55^.

Each particle projection is used as a grayscale mask which is used as per-pixel weights for simulating electron scattering events (shot noise) by sampling from a Poisson distribution inside of the mask to generate a new, unique projection. This process is divided into simulated frames (`--num_simulated_particle_frames`) to mimic an electron-counting camera, then each frame is optionally lowpass-filtered to simulate protein damage by the accumulated electron dose (`--dose_damage`, `--dose_a`, `--dose_b`, and `--dose_c`)^56^. EMAN2 is then used together with several user-specified and micrograph-specific parameters to apply CTF to each projection individually.

The AnyLabeling JSON file associated with the current micrograph is then used to optionally remove particles that are to be added near or within obscured regions from the coordinate file(s) that will be saved later (`--no_junk_filter` and `--polygon_expansion_distance`). Optionally as well, particles to be added too close to the edge of the micrograph (`--save_edge_coordinates`) and overlapping particles (`--save_overlapping_coords`) are removed from the coordinate file(s) that will be saved later. A collage of particles the size of the micrograph is then created and Gaussian noise is added to the collage based on the estimated Gaussian noise of the micrograph. The collage is added to the micrograph where the intensity of the collage’s pixels are weighted by the low-resolution features of the micrograph mimicking the ice thickness effect on particle contrast if the ‘micrograph’ `--distribution` option is selected.

Micrographs (MRC, PNG, and/or JPEG format; `--mrc`, `--png`, `--jpeg`, `--jpeg_quality`) and filtered particle coordinates (STAR, IMOD .mod, and/or COORD format; `--imod_coordinate_file`, `--coord_coordinate_file`) are then saved and are optionally downsampled (`--binning`). If IMOD is installed, the user can request that 3dmod open the newly-generated micrographs (`--view_in_3dmod`). Particles can be requested to be cropped from the micrographs, providing realistic context (`--crop_particles`, `--max_crop_particles`, `--box_size`). An information file containing relevant information for each structure set projected onto micrographs (including the structure set names, structure names, mass in kDa, the number of micrographs generated, the number of particles projected, and the number of particles saved) is saved to the run directory.

If `--parallelize_structures`, `--parallelize_micrographs`, `--gpus`, `--use_cpu`, and `--cpus` are not specified by the user, then VirtualIce will automatically determine these values to maximize the CPU and GPU usage, otherwise these user-specified settings will be respected. Multi-GPU usage is handled intelligently by distributing tasks amongst available GPUs based on GPU utilization and GPU RAM. All terminal outputs from VirtualIce may be suppressed with the `--quiet` flag so that VirtualIce may be integrated seamlessly into other workflows.

## Discussion

VirtualIce is designed to assist cryoEM analysis and software development by generating realistic, ground-truth half-synthetic micrographs where the background is real vitrified buffer and the particles mimic realistic behavior. While prior cryoEM micrograph simulation software focuses on making the forward projection model realistic at a per-particle level, VirtualIce focuses on realistic contextual particle behavior by using real background and realistic particle behavior (particle distribution based on local ice thickness, overlapping particles, particle aggregation, preferred orientation, multiple structure types per micrograph). Nevertheless, VirtualIce comprehensively models shot noise, protein dose-dependent damage, CTF, and Gaussian noise in its protein projection algorithm.

The needs of cryoEM researchers and developers is evolving as cryoEM is increasingly becoming a routine structural biology tool. Optimization and automation of every step in the cryoEM workflow will further increase the growth of cryoEM - from data collection, to real-time processing, to particle picking, alignment, and classification. VirtualIce addresses these needs by providing realistic, ground-truth cryoEM micrographs with a multitude of use-cases. Below, we expand on some possible use-cases of VirtualIce.

### Visualizing Novel Proteins in CryoEM

To help guide screening and data collection of novel proteins in cryoEM, researchers may use VirtualIce to generate micrographs with varying characteristics (e.g. preferred orientation, ice thickness, aggregation) using hypothesized models from other structure analysis techniques or speculative models from AlphaFold^57^, for example. Visual comparisons between micrographs of novel proteins in cryoEM and VirtualIce-generated micrographs may allow researchers to understand their protein’s behavior on cryoEM grids and, in turn, to better optimize their sample (e.g. reduce aggregation), plan data collection (e.g. tilted collection), and interpret their data (e.g. identify particle orientations).

### Training Particle Pickers

VirtualIce may be used to generate realistic, synthetic ground-truth datasets where only true positive particles are labeled and all true negative objects (ie. particles overlapping with other particles or obscuring objects or particles too close to the edge of the micrograph) are not labeled. The realistic background in VirtualIce datasets adds contamination and grid substrate as true negatives, which may significantly increase the precision-recall abilities of machine learning particle pickers trained on very large datasets that VirtualIce can produce by leveraging the PDB and EMDB. More accurate general particle pickers will, in turn, produce cleaner data for automated classification and alignment algorithms. VirtualIce outputs the mass of the proteins used in micrograph generation, which may then be used to train particle pickers on particles of specific sizes or types, thus creating specific general particle pickers.

### Assessing and Addressing Preferred Orientation

Preferred orientation plagues the field of cryoEM because most proteins on cryoEM grids adsorb to the air-water interfaces^55^. This systematic issue is confounded by the poor angular assignment of particles due to low signal-to-noise in cryoEM and incomplete 3D prior models^58–61^. VirtualIce-generated micrographs with ground-truth preferred orientations may be used to develop new software and refine existing software for determining angular assignments of particles.

### Creating Validation Datasets for Heterogeneity Analysis

VirtualIce allows for multiple structures to be added to micrographs. Since VirtualIce operates on atomically-defined PDB files, then ground-truth datasets may be created with as little or as much heterogeneity as desired (from single-atom/residue differences to entirely different structures). Algorithm developers interested in building software to reliably reconstruct conformationally- and compositionally-heterogeneous proteins, including continuously-heterogeneous proteins, in cryoEM can use VirtualIce as a development testbed. Benchmark datasets may be generated for community software development challenges.

### Predicting Data Collection Requirements

A key issue for determining efficient and fair distribution of cryoEM resources is in predicting how much high-resolution data collection time should be assigned for each project destined for a given microscope. However, there are no reliable ways to predict the amount of data collection time required for a given sample to reach a desired resolution, or even if that resolution is possible. VirtualIce may be used to create ground-truth datasets with varying pdb-to-mrc conversion resolutions, which may then be gradually inputted into real-time cryoEM processing pipelines, mimicking real-time collection. With this workflow, it may be possible to develop new software to determine, based on the first hours of “collection”, whether a given sample may be able to reach a desired resolution. One method for achieving this may be to continually update real-time 3D reconstructions and ResLog/Rosenthal-Henderson^62,63^ plots to determine how much data is required to predict with sufficient accuracy the actual resolution that a dataset may achieve given a specific number of more particles, and thus collection time.

### Simulating Automated Data Collection

Automated cryoEM data collection software packages are being developed with the aim of intelligently determining optimal cryoEM grids, grid squares, and holes for data acquisition^5–7,64,65^. Real-time optimization of collection areas will substantially increase collection efficiency and requires algorithms that are able to discern between high-magnification areas on the grid that meet or do not meet the structure determination needs of the biological project. VirtualIce may be used to generate ground-truth datasets for designing and training algorithms to discern between grid areas with sample issues (e.g. low particle number, preferred orientation, aggregated particles, overlapping particles, sample heterogeneity, lack of high-resolution information, low contrast) and those without significant issues. VirtualIce datasets may allow for algorithms to be designed *in-silico* orders of magnitude faster than in real collection environments because of the lack of overhead for dedicated microscope time and because of the ground-truth particle coordinate output from VirtualIce.

### Teaching CryoEM

The extensive options for producing and altering realistic particle behavior in VirtualIce provide a fruitful environment to explore for those new to cryoEM. VirtualIce may be useful as an interactive tool for understanding the extent of noise in cryoEM, particle sizes, particle orientations, preferred orientation, aggregation, particle picking, variations in defocus, and the effects of dose damage.

#### Future Developments

VirtualIce may be improved and extended in several ways. Here we describe a few potential future developments:

- 3D model of ice: VirtualIce currently does not explicitly model the vitrified buffer ice in 3-dimensions. Instead, it uses the lowest-resolution features in the vitrified buffer micrograph as a proxy for the ice thickness where darker areas are assumed to be thicker ice and lighter areas thinner ice. The defaulted micrograph distribution option in VirtualIce then uses these features as a probability map for particle placement. VirtualIce may be improved by introducing an explicit 3D ice model, possibly based on local energy-filter ice thickness^66^ estimation, which may be used to restrict particle placement based not only on particle-particle distances in the plane of the micrograph, but also in the z-direction. This development would allow for only particles and particle orientations of specific sizes to fit in thin ice, thereby improving the realism of the VirtualIce.
- Protein fragmentation: Real protein samples often have denatured or fragmented pieces of proteins in the samples, due possibly to the purification process and/or the cryoEM preparation process. These fragmentations may be modeled from PDB files inputted into VirtualIce and added to the micrographs using the existing multi-structure capabilities of VirtualIce. This development would further improve the realism of VirtualIce.
- Collect vitrified buffer micrographs from additional setups: The vitrified buffer dataset released with VirtualIce was prepared on a carbon grid and collected on a Krios with a K2 camera. Variations of grid type, microscope type (including voltage), camera type, and collection parameters will allow for more generalized use of VirtualIce-generated datasets.

We hope that VirtualIce increases development of more optimized and automated algorithms, spurs mathematical analysis and models of particles, and assists in researchers’ visual analysis of their proteins in cryoEM. VirtualIce runs efficiently on multi-CPU, multi-GPU systems, enabling realistic micrograph generation typically at a rate of 10-60 seconds per micrograph, depending on particle number and computational resources.

## Data Availability

The vitrified buffer micrographs, frames, associated labels used by VirtualIce, and green-tinted images used for manual labeling have been deposited to the public domain in the Electron Microscopy Pilot Image Archive (EMPIAR)^67^ under accession EMPIAR-12287.

## Code Availability

VirtualIce is publicly available on GitHub and Zenodo at https://github.com/alexjnoble/VirtualIce and https://doi.org/10.5281/zenodo.13852111. VirtualIce is licensed under the MIT License.

## Ethical Use Agreement

VirtualIce is for educational and research purposes, providing synthetic cryoEM data to aid in visualizing proteins in cryoEM and developing and testing image analysis algorithms. Users must clearly label any data generated with VirtualIce as synthetic to prevent deception. Users should not present synthetic data in publications or presentations without clearly labeling as synthetic so that it not be misrepresented as real experimental data.

## Acknowledgements

The author thanks Tristan Bepler (SMLC) for coming up with the idea to collect vitrified buffer micrographs for a different project and the entire SEMC group for critical assessments of this project. This work was performed at the Simons Electron Microscopy Center at the New York Structural Biology Center, with support from the Simons Foundation (SF349247) and NY State Assembly.

## Author Contributions

A.J.N. conceived of the project, wrote the software with assistance from OpenAI’s GPT, Anthropic’s Claude, and Google’s Gemini models, and wrote the manuscript.

## Competing Interests

The author declares no competing financial interests.

## Figures

**Supplementary Figure 1.**
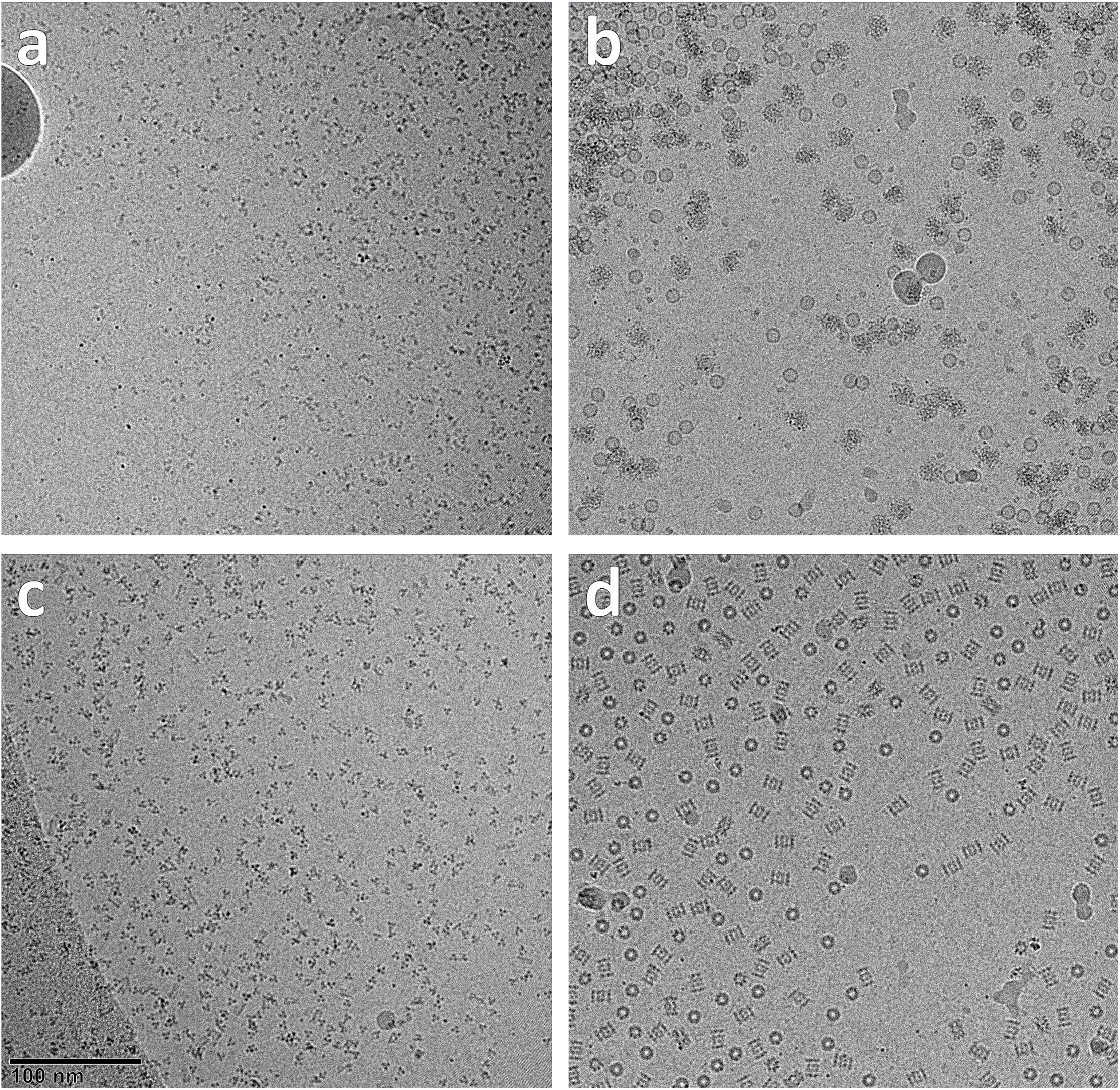
Assortment of VirtualIce-generated micrographs, unlabeled. (a) The same selection of micrographs as Figure 1 but without particle labels, illustrating the raw output of VirtualIce.

**Supplementary Figure 2.**
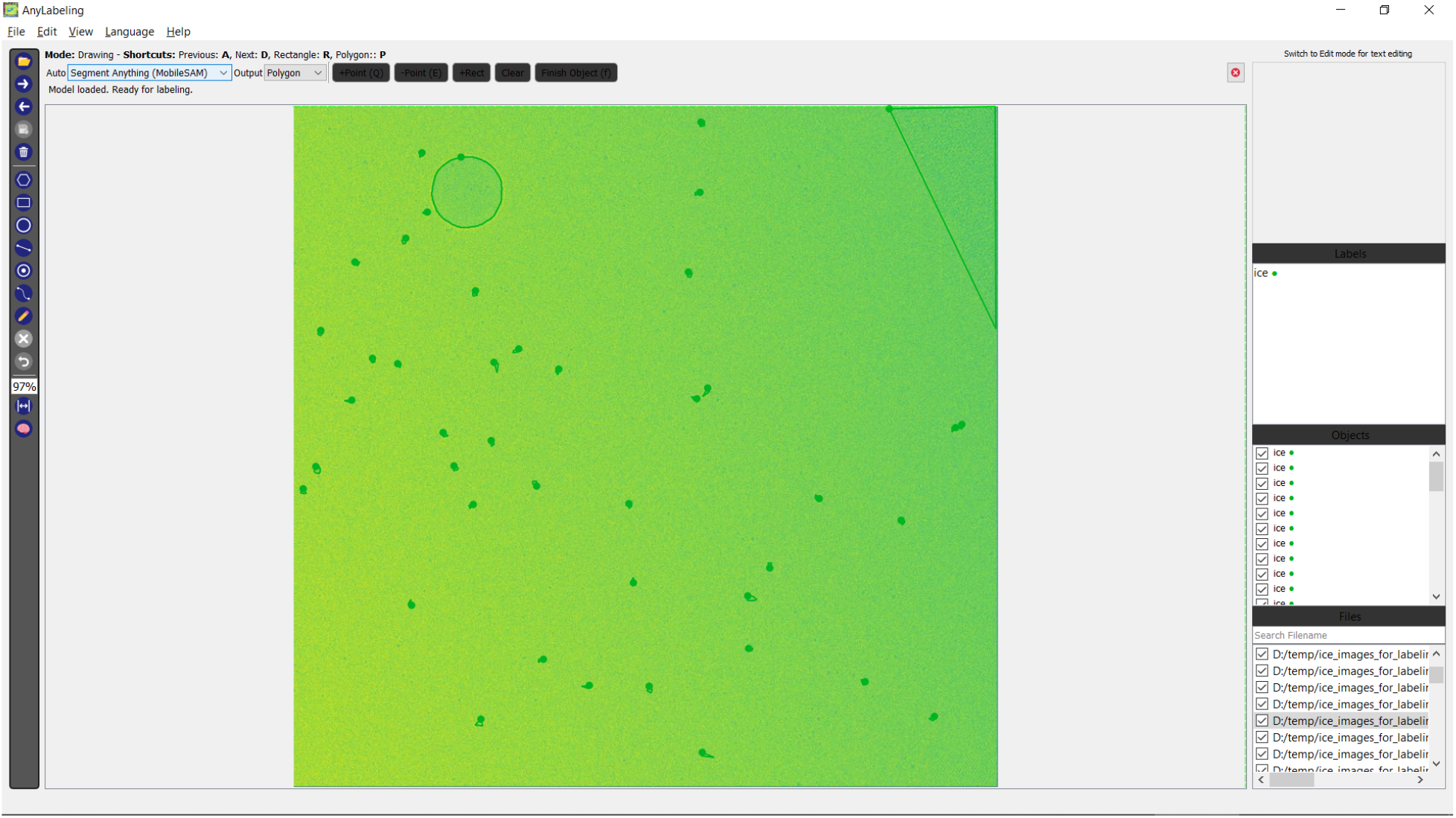
AnyLabeling software used for labeling objects that may obscure particles. Micrographs were tinted green to expedite the visualization process, then loaded into AnyLabeling where ice contamination, grid substrate, and gold flakes were manually labeled (as ‘ice’) with the assistance of MobileSAM.

## Notes

### Competing Interest Statement

The authors have declared no competing interest.

